# A truncated soil phage catechol 1,2-dioxygenase illustrates how viruses preserve and disseminate auxiliary catalytic functions in the soil microbiome

**DOI:** 10.64898/2026.03.25.714067

**Authors:** Ruonan Wu, Garry W. Buchko, John R. Cort, Deseree J. Reid, Trinidad Alfaro, Mischelle M. Schutz, Lijun Liu, Kevin P. Battaile, Scott Lovell, Ryan S. McClure, Kirsten S. Hofmockel

**Affiliations:** Earth and Biological Sciences Directorate, Pacific Northwest National Laboratory, Richland, Washington, USA; School of Molecular Biosciences, Washington State University, Pullman, Washington, USA; Seattle Structural Genomics Center for Infectious Diseases, Seattle, Washington, USA; Institute of Biological Chemistry, Washington State University, Pullman, Washington, USA; National Security Directorate, Pacific Northwest National Laboratory, Richland, Washington, USA; Protein Structure & X-Ray Crystallography Laboratory, University of Kansas, Lawrence, Kansas, USA; NYX, New York Structural Biology Center, Upton, NY, USA; Department of Agronomy, Iowa State University, Ames, Iowa, USA

**Keywords:** auxiliary metabolic gene, phage, soil, microbiome, catechol degradation

## Abstract

Bacteriophages can rewire host chemistry via auxiliary viral genes (AVGs). Using metagenomic and metatranscriptomic data from the native soil microbiome, we identified transcriptionally active AVGs, including a viral catechol 1,2-dioxygenase (V-C12DO). V-C12DO shares ∼40% sequence identity with its nearest bacterial homologs and lacks the helical dimerization domain. Despite truncation, V-C12DO retains more than ∼25% of the global consensus residues compared to C12DOs across domains of life, including the two tyrosines and two histidines that coordinate the non-heme Fe(III) active site. A 1.7 Å crystal structure also showed the conservation of the canonical β-sandwich scaffold for the iron. We next confirmed that V-C12DO cleaves catechol and remains highly active across a broad range of temperatures (30-60 °C), pH (5.5-9), and salinity (up to 2 M), exceeding those of known bacterial CD12Os. This work shows that truncated phage enzymes preserve the core catalytic chemistry and potentially further expand host metabolic versatility across dynamic environmental conditions.

## Main Text

Bacteriophages are abundant in soil, infecting a wide range of microbial hosts with diverse genetic repositories, and are increasingly recognized as integral to microbially mediated biogeochemical processes^1,2^. Phage predations on keystone microbial species can influence the microbial composition and thus the metabolic potential of soil^3,4^. In addition, the release of microbial intracellular materials after phage-induced host lysis further stimulates metabolic processes by alleviating nutrient limitations and facilitating carbon exchange within the soil microbiome^4,5^. Beyond these top-down effects, accumulating evidence suggests that phages also influence soil metabolism through expression of auxiliary viral genes (AVGs)^6,7^.

AVGs are genes that are nonessential for viral replication but are retained in viral genomes for reasons that are not fully understood. Previous research on AVGs has revealed that their abundance and functional potential can be related to environmental stress in the microbial system. For example, higher levels of AVGs related to carbohydrate metabolism were detected in more nutrient-limited subsurface soils than in surface soils^8^. Soil with higher water stress contained AVGs spanning a broader range of metabolic pathways^9^, illustrating how AVG prevalence shifts with environmental context. However, most predictions of these auxiliary metabolic functions remain sequence-based, lacking biochemical validation. Only a few soil AVGs have been verified using purified proteins expressed from the synthesized AVGs^10^, highlighting both the promise of sequence-based discovery and the major bottleneck in functional verification. Our previous study advances this field by identifying a soil AVG encoding chitosanase from forest soil metagenomes as a proof-of-concept and validating its function using enzyme activity assays, most notably by resolving an atomic-resolution crystal structure of site-specific mutants that pinpointed substrate-specific bindings^11^. These examples demonstrate that soil viruses can encode biochemically functional enzymes even though they are auxiliary to viral replication, underscoring the need for additional structure-biochemical studies to elucidate the ecological roles of environmental phages carrying diverse and active AVGs. Yet key questions remain, including whether these enzymes are expressed in situ, how robust their activities are across environmentally relevant ranges of pH, salinity, and temperature, and what ecological advantages they confer to infected hosts.

Besides carbohydrates and chitin, lignin is another plant-derived molecule that can be metabolized by soil bacteria, with catechol being a key intermediate in the lignin breakdown pathway^12–15^. The degradation of catechol is performed by two structurally and mechanistically different families of dioxygenases. Meta, or extradiol, catechol 2,3-dioxygenases (C23DO) use non-heme Fe (II) to cleave the carbon-carbon bond between the ortho hydroxyl group and the meta carbon by inserting two oxygen atoms to yield 2-hydroxymuconic semialdehyde^16,17^. In contrast, ortho, or intradiol, catechol 1,2-dioxygenases (C12DO, EC1.13.11.1) use non-heme Fe (III) to oxidatively cleave the carbon-carbon bond between the adjacent phenolic hydroxyl groups to generate cis,cis-muconic acid^18^. While the extradiol family of dioxygenases is phylogenetically ubiquitous, the intradiol family is largely confined to bacterial and fungal species^19^.

While some dioxygenase genes have been found on mobile genetic elements such as plasmids^20^, their identification in viral genomes is new, and their role in catechol degradation is unknown. For the non-plasmid and non-viral C12DO gene products, crystal structures have been determined from seven bacterial species, with one crystallizing as a monomer and six as homodimers^21–24^. The six homodimers are from *Rhodococcus opacus* 1CP (3HGI)^24^, *Pseudomonas arvilla* C-1 (2AZQ)^22^, *Acinetobacter calcoaceticus* (1DLM)^25^, *Burkholderia multivans* (9DR8), *Burkholderia vietnamiensis* (5TD3), and *Burkholderia ambifaria* (5VXT). Each of these C12DOs contains two domains, an N-terminal dimerization domain composed of three to four helices, and a C-terminal domain composed primarily of a β-sandwich scaffold for the penta-coordinated non-heme iron at the active site and an α-helix that intertwines with the helical dimerization domain. Crystal structures for two monomeric C12DOs have also been determined; one in a prokaryote: *Streptomyces* sp. SirexAA-E (4ILV)^26^ and one in a eukaryote, the spider mite, *Tetranychus urticae* Koch (6BDJ)^19^. The *Streptomyces* version lacks the N-terminal dimerization domain common to C12DOs but contains a carbohydrate-binding module (i.e., superfamily 5/12) at the C-terminal. The *Tetranychus* version also lacks the N-terminal domain common to C12DOs but instead contains an N-terminal domain of unknown function. In all C12DO crystal structures, the penta-coordinated non-heme iron is observed ligated to a water molecule and two histidine and two tyrosine side chains in a trigonal pyramidal arrangement^21–24^.

Despite growing recognition of AVGs as modulators of host metabolism, their diversity in native microbiomes, the extent to which they are expressed *in situ*, and how their molecular properties compare to their bacterial counterparts remain largely unexplored^27^. Here, we identify which AVGs are present and expressed in a Warden silt loam soil microbiome and characterize a representative AVG at the enzymatic and structural levels to directly assess how viral and bacterial homologs converge and diverge in function and architecture. This work detected a phage-encoded catechol 1,2 dioxygenase (V-C12DO) from the field site with significant sequence truncation compared to bacterial homologues, while still preserving expression and active catalytic function with robust stability across diverse soil conditions. This combined genomic, biochemical, and structural approach provides direct evidence that soil viruses can encode functional catechol-cleaving enzymes, adding to the growing list of functional AVGs. We also establish a framework for systematically validating auxiliary viral metabolic functions beyond sequence inference. These results not only reveal a soil viral innovation in catechol metabolism but also suggest a broader viral strategy to maintain auxiliary catalytic function for environmental adaptation while balancing replication cost, providing molecular insight into virus-mediated microbial processes that remain largely unexplored.

## Results

### Soil AVGs are expressed yet highly divergent from bacterial and fungal homologous genes

We observed that AVG sequences carried by soil phages exhibit substantial divergence from their bacterial and fungal homologs but may still retain functional potential. A total of 37 putative assayable AVGs assigned to 26 unique functions were previously detected from soil phage contigs recovered from replicated terabase-sized metagenomes of the Prosser, Washington grassland^7,28,29^ (**Extended Data 1**). The consistent detection of these AVGs suggests they are not transient but represent stably maintained and potentially adaptive viral functions, selected within the soil microbiome under arid grassland conditions. Although the functional annotation was previously performed using profile Hidden Markov models (HMMs) with stringent cutoffs^7^ (i.e., an E value□of <10^−15^, coverage□of > 0.35), pairwise whole sequence alignment of these AVG protein sequences against bacterial homologs in the NCBI NR database showed broad divergence, with sequence identity ranging from 27% to 77%, with an average of 48% (**Extended Data 1**). The coverage percentage of each AVG sequence aligned to the reference sequences in the NCBI NR database ranged from 25% to 99%, averaging 87%. The 37 soil AVG protein sequences generally have a high average sequence coverage, but low average sequence identity compared to the non-viral reference sequences in the public database, indicating considerable whole sequence divergence. Notably, 24 out of the 37 AVG-encoded proteins show similarity to proteins in the Protein Data Bank (PDB) and exhibit AlphaFold-predicted secondary and tertiary structure across most of their length (**Extended Data 1**). This suggests that despite sequence divergence, the AVG proteins likely retain functional folds and catalytic potential.

Many of the AVGs we detected via genomic analysis were also expressed in the paired soil metatranscriptome^30–32^, while a smaller number of cloned AVGs could be successfully expressed and soluble in *Escherichia coli*. The potential *in situ* expression of soil AVGs was conservatively assessed based on transcript reads uniquely mapped to the corresponding AVG sequences in more than one soil replicate (**Extended Data 1**). A total of 22 AVGs representing 11 unique functions were found to be potentially expressed within the studied microbiome. These expressed AVGs included glycosyl hydrolases belonging to GH5, GH19, and GH26 families, ATP-dependent Clp protease, transferases belonging to GT1 and GT4 families, asparagine synthase, acyl carrier protein, catechol 1,2 dioxygenase, and WhiB transcriptional factors. The detection of these transcripts suggests that a subset of viral auxiliary functions is under strong selective retention within the studied soil microbiome. Among the 22 AVGs with metatranscriptome-informed expression, 16 AVGs were cloned into *E. coli* expression constructs to test for *in vitro* expression. Soluble protein expression was observed for two of these AVG clones. One belonged to the GH19 family and the other was a catechol 1,2-dioxygenase. While a phage chitinase has been widely studied and experimentally validated^33^, there are no reports linking phages to catechol degradation in soil, a central step in the breakdown of lignin-derived aromatic compounds that strongly influences soil carbon turnover and organic matter formation.

### Comparative sequence analyses reveal unique sequence features for V-C12DO

The embedding-space structure of C12DOs from domains of life is primarily explained by taxonomy. Lineage accounts for roughly half of the embedding-space variance (0.498, **Fig. 1a**), whereas sequence length and broad domain categories explain much smaller fractions (0.024 and 0.012, **Fig. 1a**). Although oligomer annotations for C12DO are sparse across reference databases, the available monomer and homo 2-mer labels for this enzyme were found in multiple lineage-specific regions rather than forming a single isolated cluster (**Fig. 1a**), consistent with similar oligomeric states arising through multiple evolutionary paths.

**Figure 1.**
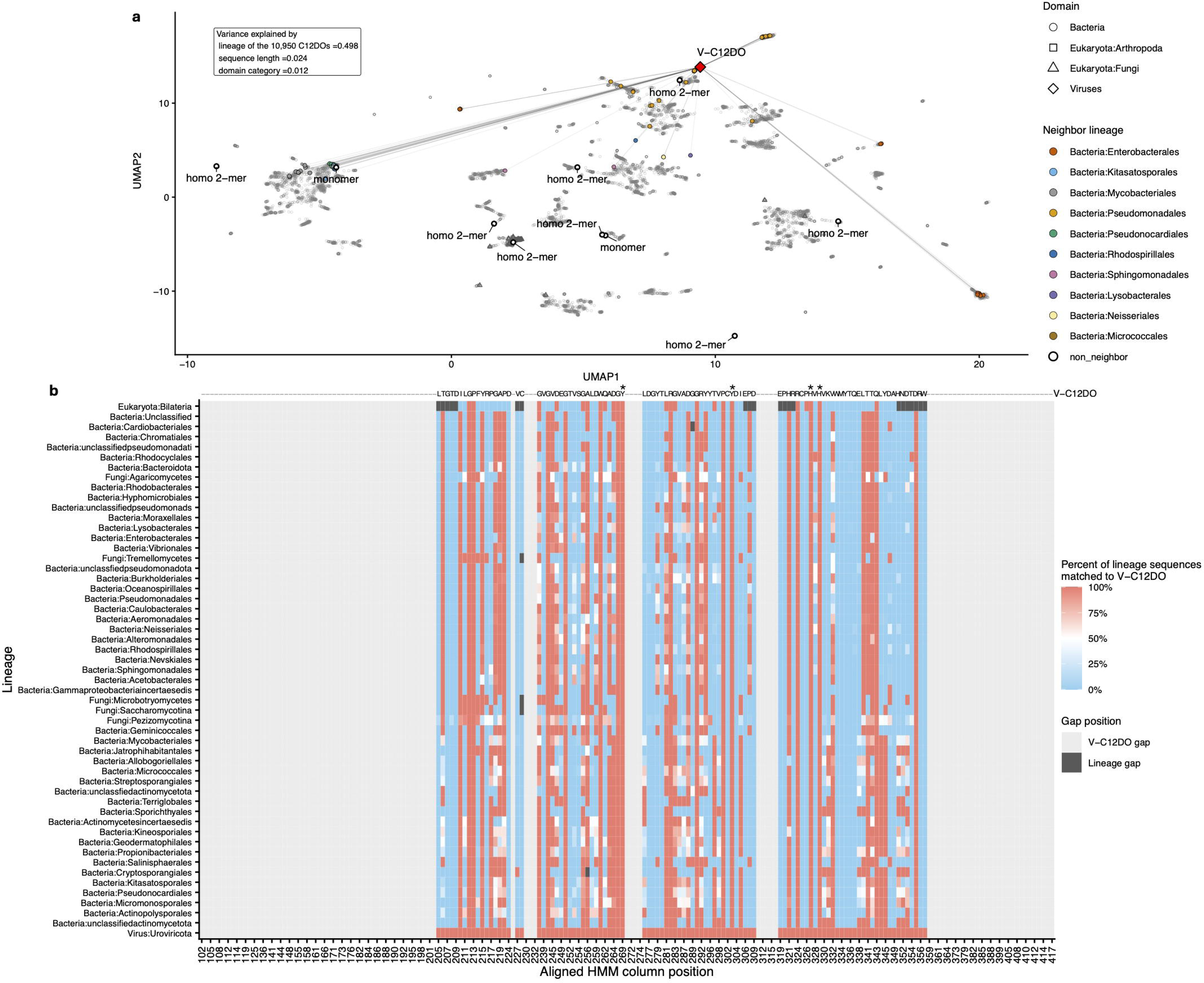
Embedding-space of C12DOs and HMM anchored residue sharing. (**a**) Uniform Manifold Approximation and Projection (UMAP) of ESM-2 sequence embeddings for 10,950 dereplicated C12DOs, with point shapes indicating domain category (Bacteria: open circles, Fungi: squares, Arthropoda: triangles, Viruses: diamonds). V-C12DO is highlighted in red, and its top 100 nearest neighbors in embedding space are colored by lineage. Database annotated oligomer states (monomer vs homo 2-mer) are overlaid as labels for the corresponding sequences. The fraction of embedding-space variance explained by lineage (partial eta squared = 0.498; Benjamini-Hochberg adjusted p value < 1 × 10□³□□), sequence length (partial eta squared = 0.024; Benjamini–Hochberg adjusted p value = 4.64 × 10□□□), and domain category (partial eta squared = 0.012; Benjamini–Hochberg adjusted p value = 8.29 × 10□³²) is reported in the inset. (**b**) HMM column-based residue comparison of V-C12DO to prokaryotic and eukaryotic lineages. For each aligned HMM column (x axis), the heatmap shows the percentage of sequences within each lineage (y axis) that match the V-C12DO residue call at that position. Only positions with strong global consensus (global maximum residue fraction >50%; the full global residue fractions are shown in Extended Data 2) are shown. Gaps are explicitly tracked, light grey indicates positions where V-C12DO has no residue call (shown as “-” in the V-C12DO track), and dark grey indicates positions where sequences in the compared lineage have no residue calls at that column. The four known key residues are marked with asterisks (*).

HMM anchoring makes deep comparisons of C12DOs interpretable despite divergence. By projecting residue calls onto aligned HMM columns and focusing on positions with strong global consensus (down-selecting global maximum residue fraction >50% from **Extended Data 2**), cross-lineage residue agreement remains well-defined even under substantial C12DO sequence divergence (**Fig. 1b**). A total of 195 HMM columns pass the >50% global-consensus filter, but V-C12DO has residue calls at only 99 columns, with 96 columns missing due to truncations and gaps (54 columns at the N-terminus, 29 at the C-terminus, 13 internal columns). Across the 99 comparable columns and 53 lineages, lineage-specific gaps are rare, with only 26 of 5,247 lineage-by-column cells missing (0.5%). A subset of positions shows broad cross-lineage agreement with V-C12DO, including 23 columns where more than 75% of lineages match V-C12DO at ≥90% frequency. Two columns are fully conserved across all 53 lineages, Position 324 (R) and Position 329 (H), and one column is fully conserved across all lineages with residue calls at that position, Position 355 (D) (among 52 lineages with calls). The four key sites (2Y and 2H, asterisks in **Fig. 1b**) map to HMM positions 269/303 (Y) and 327/329 (H) and are among the most conserved positions overall, with Position 329 fully conserved (53/53 at 100% match) and the remaining three sites showing near-complete agreement (minimum match fractions of 99.3-99.9%, with 50-52 lineages at 100% depending on the site). This HMM-anchored global sequence mapping enables direct comparisons between V-C12DO and diverse prokaryotic and eukaryotic C12DOs without relying on high overall sequence identity or comparable sequence length, while still resolving residue-level conservation and divergence. The conservation of four key residues (two tyrosine and two histidine) that coordinate iron in the active site, the shortness of the V-C12DO protein sequence relative to C12DOs in bacterial clades, and our ability to successfully express and purify soluble protein motivated further investigation of the biochemical and structural properties of V-C12DO.

### Enzymatic assay of V-C12DO confirms substrate specificity and enhanced adaptability

V-C12DO catalyzes the same reaction as bacterial catechol 1,2-dioxygenases but exhibits greater tolerance to pH, salinity, and temperature, indicating enhanced adaptability of this AVG-encoded enzyme. When assayed with catechol as the substrate, V-C12DO displayed a characteristic absorption increase at 260 nm over time at room temperature (20 °C) and pH 6.5 (its optimal pH shown below), while no increase at 375 nm was detected under any experimental condition (**Fig. 2a**). This pattern confirms that V-C12DO functions as a catechol 1,2-dioxygenase rather than a catechol 2,3-dioxygenase, whose product absorbs at 375 nm^34,35^. The measured K_m_ and V_max_ of V-C12DO for catechol were 43.5 μM and 27.9 μM/min, respectively (**Fig. 2b**), the former value about one order of magnitude higher than measurements made for other C12DOs^36^, suggesting that it may be modestly less active.

**Figure 2.**
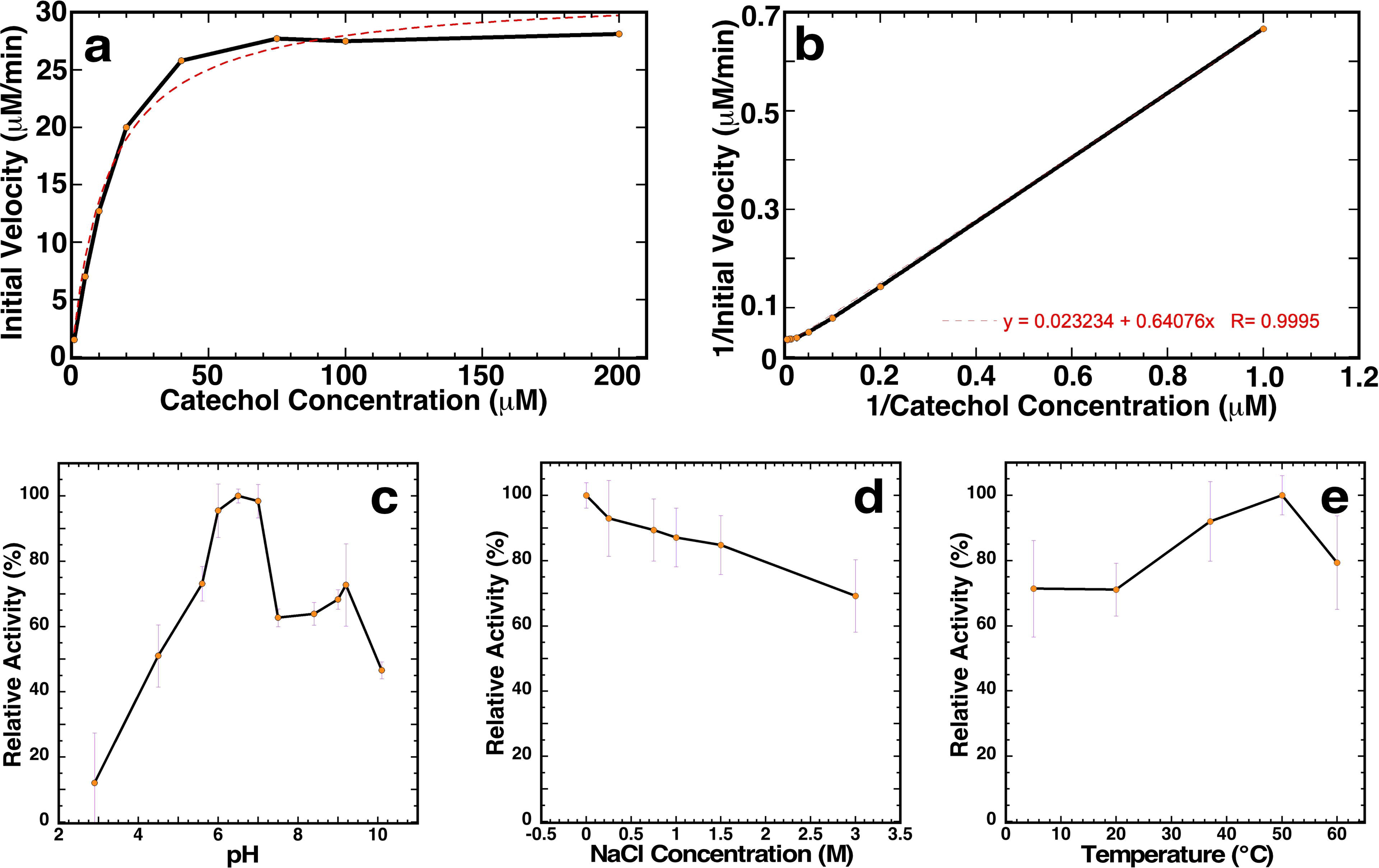
Effect of catechol concentration, pH, salinity, and temperature on the enzymatic activity of V-C12DO. The enzymatic assays were performed using the standard reaction buffer (50 mM sodium phosphate, 100 mM NaCl, pH 6.5). (**a**) Michaelis-Menton plot of the experimental data collected at 20 °C. Each point is the mean of three experiments, with the error bars reflecting the standard deviation. (**b**) A double-reciprocal Lineweaver-Burk plot of the experimental data in (**a**). The y-intercept is 1/V_max_, V_max_ = 43.5 μM/min. The slope is K_m_/V_max_, therefore K_m_ = 27.9 μM. **(c)** Summary of the pH activity assays performed at 20 °C in 100 mM NaCl and 50 mM sodium acetate (pH 2.8 – 5.7), 50 mM BisTris (pH 6.0), 50 mM sodium phosphate (pH 6.5 – 7.5), 50 mM Tris (pH 8.4 – 9.0), and 50 mM CAPS (pH 10.1). (**d**) Summary of the sodium chloride activity assays performed at 20 °C in 50 mM sodium phosphate buffer (pH 6.5) with NaCl concentrations ranging between 0.1 and 3.0 M. (**f**) Summary of the thermal stability activity assays performed in the standard reaction buffer between 5-60 °C. In assays C through E, each point is the mean of three experiments, with the error bars reflecting the standard deviation.

V-C12DO is specific for catechol while displaying only limited tolerance for substituted analogs. We found that substitution of the target molecule at the 3-position with a methyl group substantially reduced activity by ∼40%, whereas substitution at the 4-position results in different enzymatic activities depending on the type of ring substituent (**Table 1**). A 4-methyl group enhanced activity to ∼166%, indicating that the enzyme can accommodate, and even favor, certain para substitutions. Similarly, a 4-ethyl group maintained activity at a level comparable to catechol (∼102%). In contrast, a 4-chloro substituent strongly impaired catalysis (11.4%), and the dichlorinated derivative 4,5-dichlorocatechol showed no detectable activity at all. Pyrogallol, which introduces an additional hydroxyl group, was also considered as an off-target substrate (2.3%). These results demonstrate that V-C12DO requires an intact 1,2-dihydroxyl motif and that para substitutions strongly influence catalytic efficiency, enhanced by small alkyl groups but suppressed by chlorine atoms. Although the list of substituted analogs examined here is not exhaustive, the same patterns of variation in substrate specificity are consistently reported for other bacterial C12DOs^24,37,38^, because the electronic environment around the active-site iron modulates reactivity^39^. These data show that V-C12DO displays activity against a range of substituted catechols, which is consistent with what has been observed for known bacterial catechol dioxygenases^40,41^.

**Table 1.** Substrate specificity of V-C12DO.

| Compound | Relative activity (%) |
| --- | --- |
| catechol | 100 ± 0.8 |
| 3-methylcatechol | 41 ± 3 |
| 4-methylcatechol | 166 ± 5 |
| 4-ethylcatechol | 102 ± 5 |
| 4-chlorocatechol | 11.4 ± 0.8 |
| 4,5-dichlorocatechol | n.d. |
| pyrogallol | 2.3 ± 0.6 |

Beyond confirming activity under standard assay conditions, V-C12DO was found to be active in a broader range of pH, salinity, and temperature compared to typical catechol 1,2-dioxygenases, reflecting potentially enhanced environmental resilience. V-C12DO exhibited maximal activity near neutral pH (∼7) and maintained more than 60% of its maximal activity across a broad range from pH 6 to 9 (**Fig. 2c**), indicating an unusually wide activity range compared with the basic optima typical of bacterial C12DOs^19,24,42,43^. V-C12DO also shows more tolerance to salinity than other assayed C12DOs, maintaining ∼90% activity at 1 M NaCl and losing only about 30% of its activity at a salt concentration of 3 M (**Fig. 2d**). The optimal temperature for V-C12DO activity is ∼50 °C (**Fig. 2e**), a value slightly above the maximum range observed for other C12DOs (30-45 °C)^38,42,44^. These results indicate that V-C12DO can maintain catalytic efficiency across broad physicochemical conditions, more so than bacterial counterparts, underscoring its structural stability that may be favorable for environmental adaptability.

### V-C12DO crystal structure reveals a conserved core with unique pentacoordinated iron

The crystal structure of V-C12DO revealed a compact fold and an unconventional metal coordination at the active-site. After extensive construct screening, we crystallized this enzyme (termed V-C12DO*) in space group P2_1_ 2_1_ 2_1_ with two chains per asymmetric unit (**Fig. 3a**; for detailed V-C12DO* crystallographic data, refer to **Extended Data 3**). Continuous electron density was observed except for the first three residues at the N-terminus and the last eight and six residues at the C-terminus in both chains. Overall, the crystal structure revealed a core, compact, all β-strand fold with metal coordination at the active site that anchors a unique dimeric interface. At the catalytic center, each chain contained a non-heme iron atom coordinated intramolecularly to two histidine residues (H118 and H120) and two tyrosine residues (Y72 and Y101), and intermolecularly to one tyrosine residue (Y5) from the opposite chain, forming a third iron-tyrosine coordination bond (**Fig. 3b**). Such an intermolecular iron-tyrosine coordination has never been observed in bacterial C12DOs, where this position is usually occupied by a water or hydroxide ligand and occasionally a benzoate-like ion^24^. This iron-tyrosine interaction extended across an 18-residue interface (Y4-P22). PDBe PISA analysis predicts V-C12DO may be a dimer in solution; however, the Complex Formation Significance Score is low (CSS = 0.328), suggesting the predicted dimer may be due to crystal packing forces rather than true physiological interactions.

**Figure 3.**
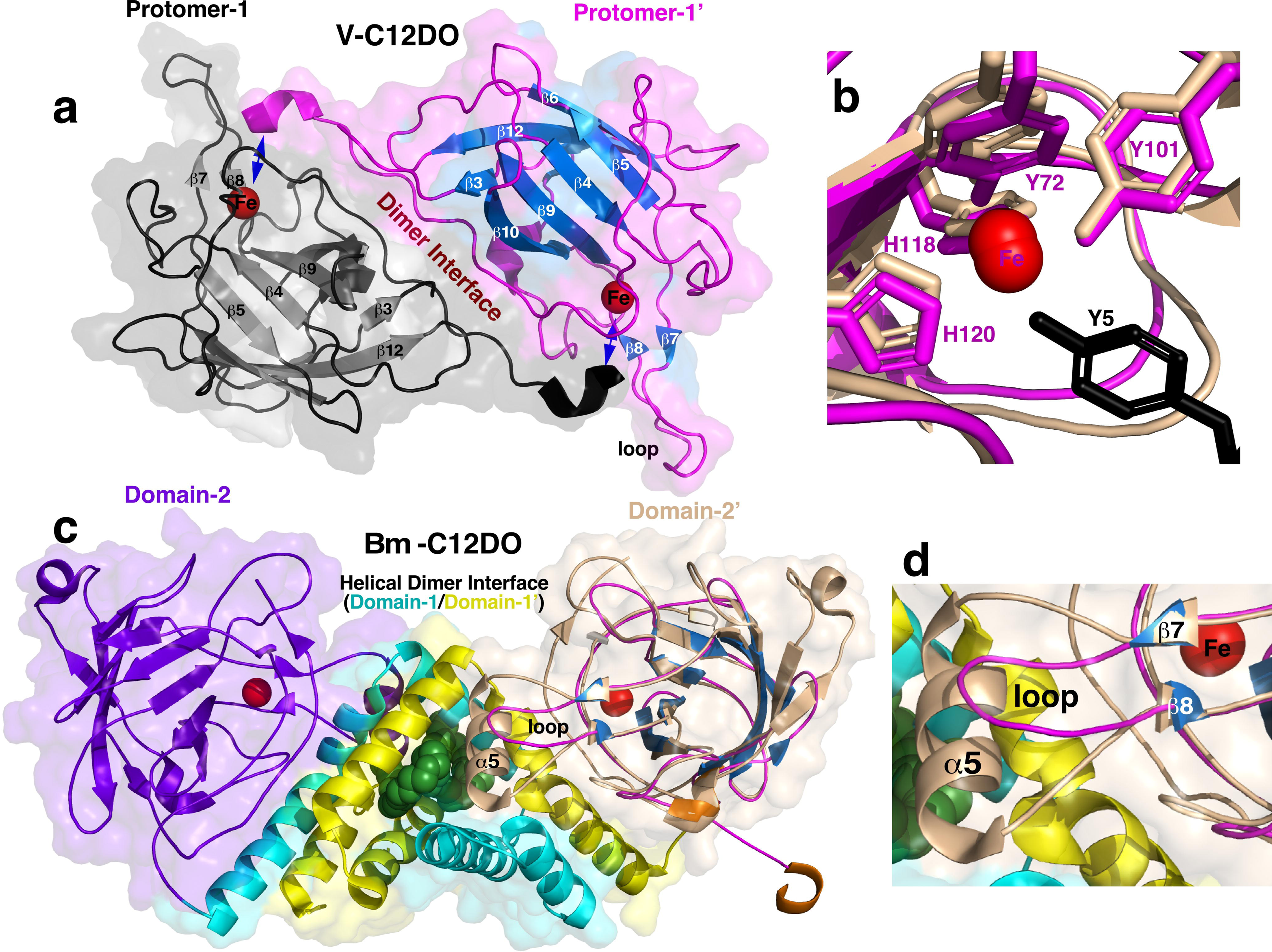
Crystal structure of V-C12DO* and comparison to a representative bacterial C12DO. (**a**) Cartoon representation of the crystal structure for V-C12DO (9YH9), a dimer with the β-strands colored gray and blue in Protomer-1 (black) and Protomer-1’ (magenta), respectively, and the iron atom in each protomer illustrated as red spheres. The dimer interface is composed of an extended unstructured region between G4-G21, held together by 18 hydrogen bonds (side chain – backbone and side chain – side chain) and anchored at each end with metal-hydroxyl intermolecular interactions between the side chain of Y5 and the non-heme iron (blue double-arrowed line, see (**b**) for expansion). (**b-d**) Various views of the overlayed structure of V-C12DO* Protomer-1’onto the dimeric structure of *Bm*-C12DO (9DR8) using the RCSB PDB online structure alignment program (https://www.rcsb.org/alignment). The following coloring schemes are used in these cartoon representations: V-C12DO: Protomer-1(black, not shown in **c** or **d**), Protomer-1’ (wheat, with β-strands colored blue in **c**). *Bm*-C12DO: Domain-1 (cyan), Domain-2 (purple), Domain-1’ (yellow), Domain-2’ (wheat), organic component (spheres colored green), and iron (red spheres). (**b**) The side chains coordinated to the non-heme iron at the active site overlay well for V-C12DO* (magenta) and *Bm*-C12DO (wheat), indirectly supporting a similar catechol 1,2-dioxygenase activity for both proteins. The major difference at both active sites is the coordination of the hydroxyl group of a third tyrosine (Y5, black) to the iron in V-C12DO* from the second protein in the dimer. (**c**) The structure of *Bm*-C12DO resembles a boomerang with the C-terminal, independent, catalytic domains (purple and wheat) at the ends held together by the N-terminal, all helical domains (cyan and yellow) in the middle. As with all crystal structures of dimeric bacterial C12DOs, phospholipid (green spheres) is present in the hydrophobic core of the helical dimer interface. The superposition of the structures of V-C12DO* Protomer-1’ (magenta with blue β-strands) onto the structure of *Bm*-C12DO domain-2’ (wheat) shows that similar folds (backbone RMSD 2.67 Å) hold the active site metal. (**d**) Expansion of the region between the conserved, sequential tyrosine (V-C12DO*: Y72) and histidine (V-C12DO*: H120) residues coordinated to the iron shows that the fifth helix (α5) in *Bm*-C12DO that interacts with the helical dimer interface is replaced by a seven-residue shorter sequence that forms an unstructured loop in V-C12DO*. All secondary structure labelling is with reference to the numbering scheme for dimeric C12DOs presented in Figure 3.

Our comparative structural analyses show V-C12DO with the canonical C12DO fold family but has a streamlined structure with key oligomerization elements removed. Beyond the N-terminal interface, V-C12DO* contains a core structure consisting of a four-strand anti-parallel β sheet stacked against a four-strand mixed β sheet (with one strand containing a two-residue β bulge) to form a β sandwich (**Fig. 3a** and **Fig. 4a**). A DALI search^45^ of V-C12DO structures (PDB ID: 9YH9) identified four bacterial catechol 1,2-dioxygenases with iron in the active site as the closest hits (Z-scores > 18.0; 3HGI, 2AZQ, 9DR8, 1DLM). The core secondary structure of V-C12DO aligns closely with the four bacterial homologs, except for the absence of two short N-terminal β-strands (β1, β2) and helix α5 (**Fig. 3a** and **Fig. 4a**). It should be noted that many other structures also returned Z-scores greater than 18 but were annotated as belonging to the other families of intradiol-cleaving enzymes. Structural superposition of V-C12DO Protomer-1 with a representative bacterial C12DO homologs (*Bm*-C12DO) shows that the viral enzyme (1) retains a canonical C12DO-like β-sandwich core composed of two stacked β-sheets, one antiparallel and one mixed, that houses the catalytic iron (**Fig. 3a** and **Fig. 3c**), (2) lacks the N-terminal four-helix dimerization domain (Domain-1, **Fig. 3c**) and (3) replaces α5 with a shorter, unstructured loop between β7 and β8 (**Fig. 3d**). The absence of the four-helix dimerization domain and α5, which form the dimer interface in bacterial C12DOs, strongly suggests that V-C12DO functions primarily as a monomer in solution.

**Figure 4.**
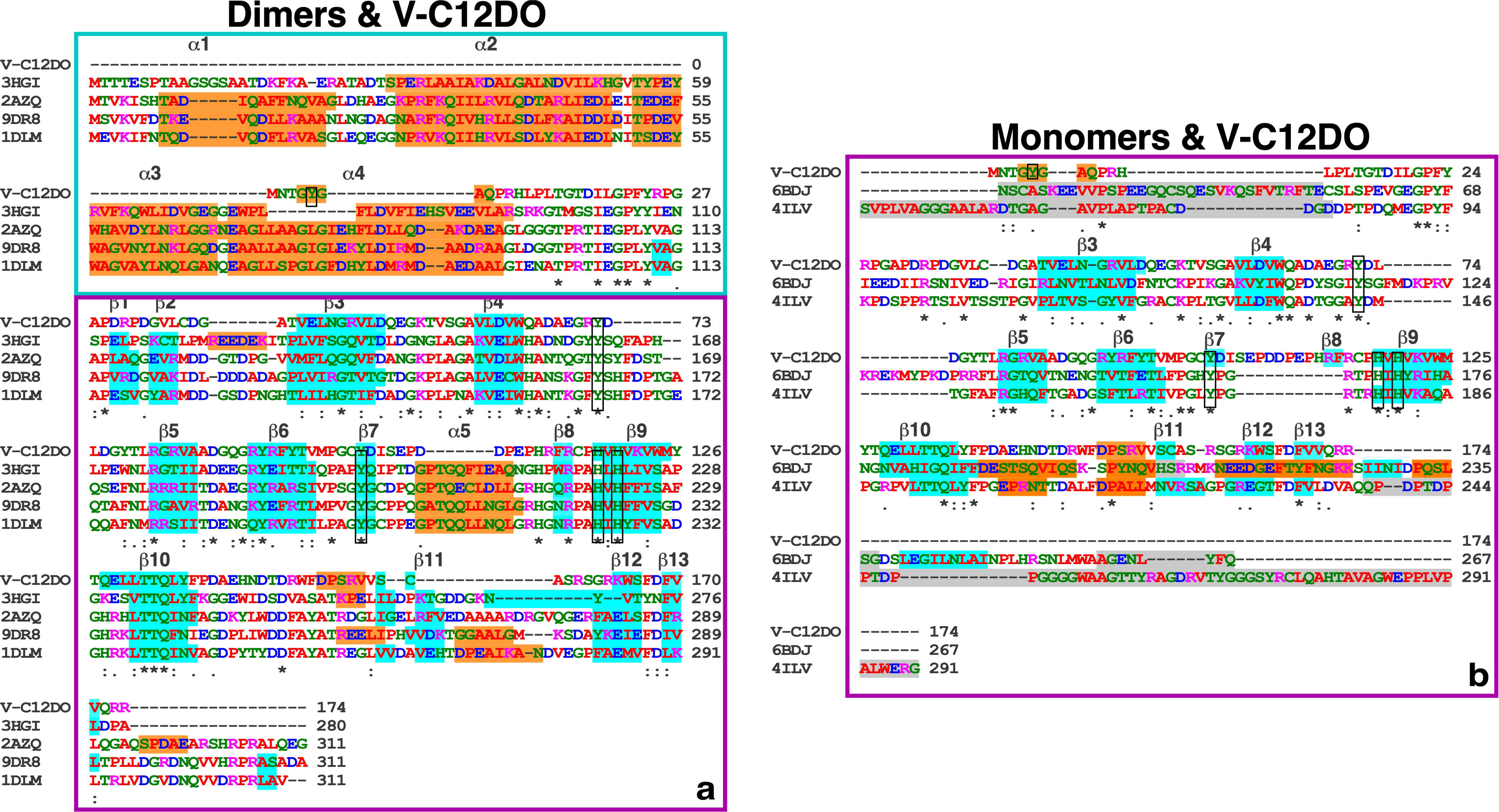
Multiple sequence alignment of V-C12DO with the sequences of C12DOs with crystal structures. (**a**) Clustal Omega multiple sequence alignment of V-C12DO (9YH9) with four bacterial C12DOs whose crystal structures were solved in the iron-bound state: 3HGI (*Rhodococcus opacus* 1CP), 2AZQ (*Pseudomonas arvilla* C1), 9DR8 (*Burkholderia multivorans*), and 1DLM (*Acinetobacter calcoaceticus*). Secondary-structure elements from the structures are overlaid, with α-helices shaded orange and β-strands shaded cyan. The two Tyr and two His residues coordinating the catalytic iron are boxed in black, together with the additional iron-coordinating residue unique to V-C12DO (Y5). Domain 1 (N-terminal dimerization domain plus linker) is boxed in cyan, and Domain 2 (catalytic core containing the active site) is boxed in purple. Helices (α1–α5) and strands (β1–β13) shared across the bacterial structures are numbered above the alignment. Residues are colored by the ClustalW scheme, and conservation is indicated below (asterisk, identical; colon, conserved; period, semi-conserved). (**b**) Clustal Omega alignment of V-C12DO with two monomeric C12DOs whose crystal structures were solved in the iron-bound state: 6BDJ (*Tetranychus urticae* Koch) and 4ILV (*Streptomyces sp.* SirexAA-E). Secondary structure is overlaid as in (**a**) with regions lacking electron density shaded grey. The iron-coordinating tyrosine and histidine residues and V-C12DO Y5 are boxed in black. The β-strands in V-C12DO are numbered above the alignment using the bacterial dimer labeling scheme in (**a**). Residue coloring and conservation symbols follow (**a**).

Sequence and structural comparisons showed that dimer formation is not required for C12DO catalytic activity, and this is further corroborated by the sequence and structures of other known, monomeric C12DOs. Two monomeric C12DOs, one from a cellulolytic bacterium (4ILV) and another from a polyphagous spider mite (6BDJ), closely align with V-C12DO with DALI Z-scores above 20 and backbone RMSDs ≤ 1.7 Å. All three enzymes share the same β-sandwich catalytic core and conserve the residues that coordinate the metal center (**Fig. 4b** and **Fig. 5a**), supporting a common catalytic mechanism. However, their N-terminal regions differ. Similar to V-C12DO, both monomeric structures lack the N-terminal helical dimerization domain found in dimeric bacterial C12DO, with no electron density or defined secondary structure present before the first β-strand of the β-sandwich core. Different from the unstructured loop between β7 and β8 seen in V-C12DO, the two monomeric homologs have an 11-residue-shorter segment here that forms a short α-helix (α5) in dimeric C12DOs that helps stabilize the helical dimeric interface (**Fig. 4b** and **Fig. 5b**). These comparisons show that V-C12DO retains the conserved β-sandwich catalytic framework with a truncated architecture that distinguishes it from both classical dimeric and monomeric C12DOs.

**Figure 5.**
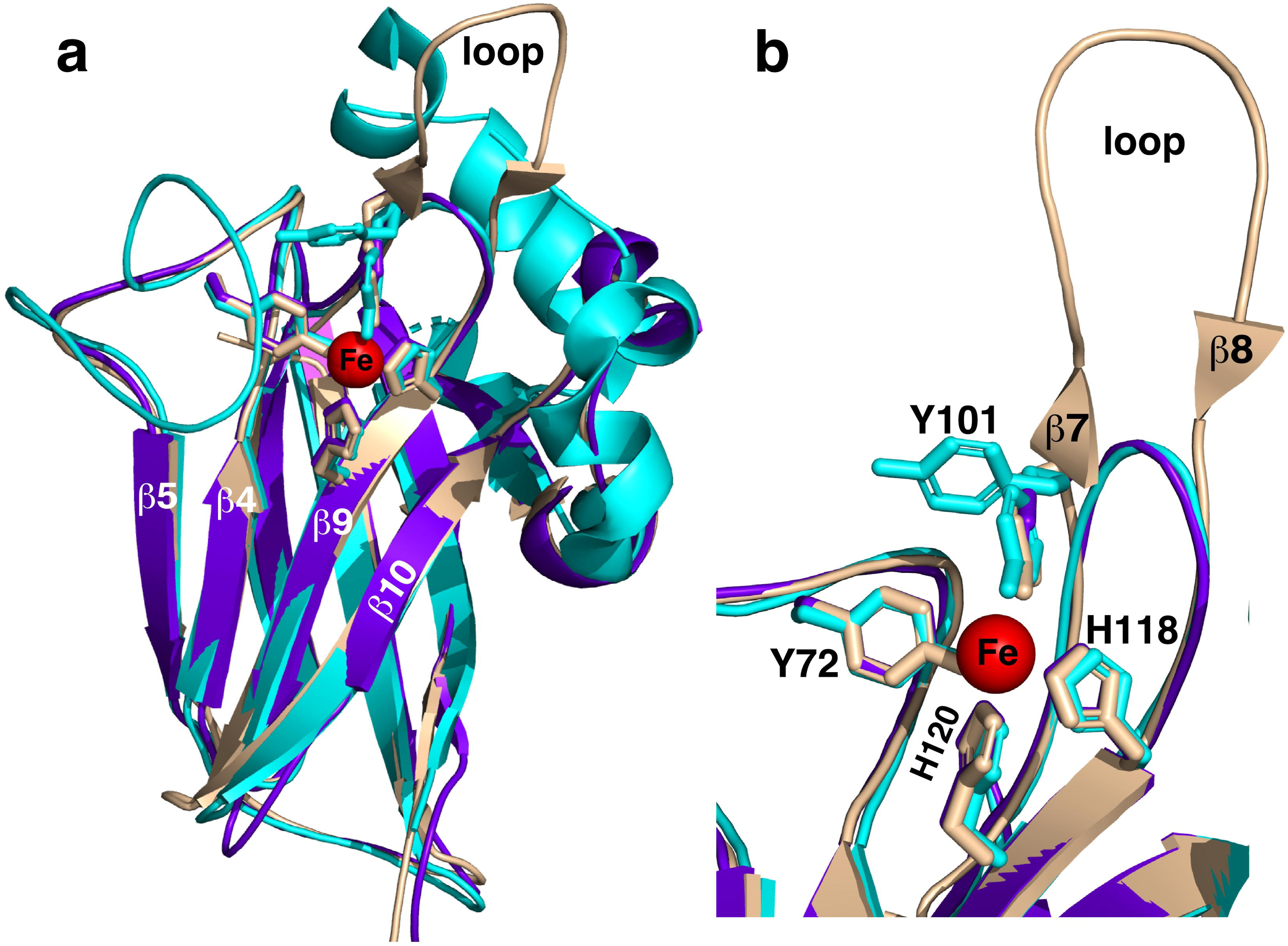
Comparison of the crystal structure of V-C12DO with two monomeric C12DO crystal structures. (**a**) Aligned cartoon representations of the V-C12DO Protomer-1’ structure (9YH9, wheat) onto the crystal structure for the monomeric C12DOs from *Tetranychus urticae* (6BDJ; cyan) and *Streptomyces sp.* SirexAA-E (4ILV; purple). Disordered regions at the termini of the structures have been removed for clarity. In both crystal structures, a non-heme iron (red sphere) is ligated to two histidine and two tyrosine side chains (illustrated as sticks). All three structures share the β-sandwich scaffold that holds the active site. (**b**) An expansion of the active site highlighting the differences in the region containing β7 and β8 in V-C12DO, with background structure removed for clarity. These two β-strands are absent in both monomeric C12DO structures, and the loop is much shorter. In the dimeric C12DO crystal structures (Fig. 3d), an α-helix exists between β7 and β8 that forms part of the helical dimer interface. The labeling scheme is as described in Figure 4a.

### Multiple techniques show conflicting evidence for the quaternary state of V-C12DO

To determine the quaternary structure in solution, V-C12DO* and the dimeric bacterial C12DO *Bm*-C12DO (9DR8) were further analyzed by complementary biophysical methods. Size-exclusion chromatography showed that V-C12DO* eluted at a retention time more consistent with the molecular weight of a ∼44 kDa dimer rather than for a ∼22 kDa monomer, eluting later than, but near, the ∼70 kDa *Bm*-C12DO dimer (**Supplementary Figure 1**). Native mass spectrometry in 200 mM ammonium acetate (pH 6.4) confirmed a dimeric species for *Bm*-C12DO but revealed the predominant V-C12DO* species as monomeric (**Supplementary Figure 2**). The ^15^N-NMR relaxation experiments yielded rotational correlation times (τ_c_) of 8.6 ± 1.4 ns for V-C12DO* and 16.6 ± 3.6 ns for *Bm*-C12DO, consistent with monomeric and dimeric states, respectively^46^. The crystallography (∼7 mg/mL; dimer), size exclusion chromatography (∼2 mg/mL; dimer), mass spectrometry (∼ 0.2 mg/mL; monomer), and NMR spectroscopy (∼10 mg/mL: monomer) gave no definitive conclusion as to whether V-C12DO is predominantly monomeric or dimeric in solution.

## Discussion

### Functional retention under genomic constraint

The persistence of AVGs in natural systems reflects selection to retain enzymes that carry essential biochemical functions while remaining structurally and functionally flexible enough to accommodate the pressure for genome streaming in phages. The AVGs identified in the studied soils show extensive sequence divergence from bacterial and fungal homologs. These soil AVGs, however, retain the conserved residues of the respective protein families to maintain the structural integrity necessary for their core functions while reducing sequence complexity. This pattern suggests that some of the AVGs are not random remnants of host DNA, but functional variants shaped by viral-specific constraints. This is consistent with what has been observed in marine viral genomes, where AVGs maintain the core catalytic functions of the system while adapting to viral replication strategies^47^, especially for those related to marine photosystems^48^. A similar strategy is also reported in recent soil carbon cycling-related AVGs^11^. Such selective maintenance of minimal yet functional gene architecture reflects adaptation to the constraints of viral genomic replication during infection. Transcriptional evidence at the microbiome level in this study further supports the functional persistence of these viral genes. The expression of viral metabolic enzymes during infection suggests that viruses can augment host metabolism to enhance viral replication, especially under nutrient-limited or stress conditions common in the studied arid soils^49,50^. Together, these results suggest that AVGs evolve to preserve catalytic function while minimizing genetic cost, allowing viruses to maintain auxiliary metabolic potential without compromising replication efficiency.

### Evolutionary divergence and convergence in C12DOs

Among the soil AVGs identified, V-C12DO provides a mechanistic example of how phage-encoded enzymes preserve function while adapting to genomic and environmental constraints. Our embedding and HMM-anchored analyses support a structural model with a conserved core and a flexible periphery. The global sequence space is strongly structured by lineage, but the residue-level constraints are governed by a small set of positions that remain conserved across distant lineages even when surrounding regions diverge. V-C12DO retains catalytic capability despite extensive sequence divergence and domain loss when compared to the non-viral C12DOs across domains of life. Although overall it shares only ∼40% sequence identity with the closest bacterial C12DOs, it conserves the active-site residues and catalyzes the same catechol ring-cleaving reaction. This pattern is consistent with other AVGs, in which catalytic cores are maintained, but neighboring regions diversify under pressure for compact genomes, which could be advantageous during replication^6,11,51,52^.

In addition to the sequence comparison, structural data further support this interpretation. V-C12DO lacks the N-terminal helical dimerization domain found in bacterial homologs, and instead, a dimer is observed in the crystal structure stabilized through an unusual intermolecular iron-tyrosine coordination. The absence of the helical dimerization domain, which in bacteria is hypothesized to help regulate product turnover^25^, suggests that V-C12DO may function independently from host regulatory control once expressed. This configuration further enables a monomeric and self-sufficient enzyme without loss of activity. Comparative structural analyses with two known monomeric C12DOs from *Streptomyces* sp. SirexAA-E (4ILV), a cellulolytic bacterium^26^, and *Tetranychus urticae* Koch (6BDJ), a spider mite^19^, show strong structural convergence. These monomeric C12DOs lack the dimerization domain and phospholipid-binding site typical of classical dimeric bacterial C12DOs^22^. Such convergence toward compact, autonomous catalytic units reflects a broader viral strategy in which structural simplification enhances replication efficiency and metabolic flexibility. Note that in this convergence to structural simplicity, V-C12DO, like the monomeric C12DOs, still has an N-terminal extension that may be necessary to prevent edge-to-edge aggregation of β-sheets^53–55^ (**Supplementary Figure 3**).

Despite domain loss, V-C12DO maintains high activity across broad pH, salinity, and temperature ranges, showing that viral genome streamlining does not compromise enzyme stability or catalytic performance. These findings suggest that phage evolution balances sequence divergence and structural convergence to preserve enzymatic capability at low genetic cost while also adapting to a wide range of environmental conditions. V-C12DO exemplifies an efficient, environmentally resilient biocatalyst while maintaining minimal genomic burden.

It should be noted that we propose the hypothesis that V-C12DO lacks much of the structure of bacterial C12DOs under evolutionary pressure, but other explanations are possible. Monomeric C12DO’s do exist: *Streptomyces* sp. SirexAA-E (4ILV), a cellulolytic bacterium^26^, and *Tetranychus urticae* Koch (6BDJ), a plant-eating spider mite^19^. Perhaps it is not a coincidence that both organisms break down cellulose and their monomeric C12DOs assist in this process. If so, V-C12DO may serve a similar role in the soil microbiome. These two non-viral monomeric C12DOs lack the dimerization domain and phospholipid-binding site typical of classical dimeric bacterial C12DOs^22^. While our findings strongly support the hypothesis that V-C12DO evolved under viral pressure to streamline genomes, alternative hypotheses, such as its derivation from existing compact microbial enzymes, remain plausible but lack experimental validation. Given the structural and functional data presented, evolutionary pressure for compatible genomes likely represents the primary driver of V-C12DO adaptation.

### Biotechnological potential of V-C12DO

The discovery and characterization of V-C12DO highlight new opportunities for developing sustainable approaches to adipic acid production. This is because the product of the oxidative cleavage of catechol, cis,cis-muconic acid, may be readily converted into adipic acid ((CH_2_)_4_(COOH)_2_) via hydrogenation. Adipic acid is a six-carbon dicarboxylic acid and a precursor for nylon, a growing, $30+ bn, global industry^56^. It is commercially obtained via an environmentally unfriendly petrochemical route^57^. A more environmentally friendly route would be to produce cis,cis-muconic acid via fermentation, followed by hydrogenation into adipic acid^58^. While a number of bacteria and yeast containing C12DO genes produce cis,cis-muconic acid via the catabolism of aromatic compounds, no known organism endogenously produces cis,cis-muconic acid from renewable sugars. However, through the introduction of heterologous synthetic pathways that exploit naturally occurring intermediates in the shikimate pathway, it is possible to produce cis,cis-muconic acid from glucose^58,59^. Because the conversion of catechol to cis,cis-muconic acid by C12DO is the final step in such schemes, productivity may be significantly improved with a C12DO of higher activity. Hence, the catalytic properties and substrate promiscuity for many C12DO have been comprehensively described^24,38,43,44,60^, providing a strong rationale for similarly characterizing V-C12DO here.

To address this pressing need, viral C12DO with less structural complexity than those found in bacteria could be promising candidates. Such compact viral enzymes can have practical advantages, including more efficient expression and folding in heterologous hosts and greater tolerance to diverse environmental conditions^61^. Their simpler structures also tend to be more accessible for modification, allowing targeted improvements in activity or substrate range^62^. These properties, shaped by viral evolutionary pressure for efficiency and adaptability, indicate the potential of V-C12DO as biocatalysts for sustainable adipic acid production. Additional AVGs found in future studies, especially those with divergent structures compared to known bacterial counterparts, could also act as new target enzymes for bioeconomy goals.

### Ecological implications of AVGs

The discovery of a functional catechol 1,2-dioxygenase encoded by a soil virus with a simplified protein structure represents a significant advancement in understanding the ecological roles of viruses within soil ecosystems. Traditionally, the degradation of aromatic compounds, essential for carbon cycling and the detoxification of organic pollutants, has been attributed to soil bacteria and fungi alone^63^. Our findings challenge that paradigm by suggesting that soil viruses may actively contribute to these processes through expressing functional AVGs, such as those encoding catechol 1,2-dioxygenase. The simplified structure of the viral enzyme may reflect adaptations to the constraints of viral-host interactions, enabling efficient expression and metabolic enhancement during infection. These adaptations could provide valuable insights into novel biochemical pathways for breaking down recalcitrant organic molecules, particularly under conditions where microbial metabolism alone is insufficient^11,42,64^. Additionally, this discovery emphasizes the underappreciated role of viruses in facilitating metabolic innovation and highlights their potential influence on the functional diversity and resilience of soil microbial communities. The potential for viruses to actively degrade pollutants through expressed AVGs is intriguing, but more analysis of quantitative ecological data is needed to distinguish viral contributions in this area relative to those of bacteria or fungi.

By encoding enzymes that support the breakdown of complex organic molecules, soil viruses may accelerate the turnover of organic matter, enhancing nutrient availability and stabilizing microbial communities^1^. Their ability to degrade pollutants, such as those from agricultural and industrial sources, also positions soil viruses as potential players in natural bioremediation processes. The presence of functional metabolic genes in viruses raises critical questions about the extent of viral influence on biogeochemical processes and microbial dynamics^65^. Future research may prioritize characterizing and quantifying the function of viral-encoded enzymes in environments under varying conditions to fully understand their ecological impact at the community and ecosystem levels. This study not only expands the understanding of the roles of soil viruses but also highlights their potential as tools for addressing ecological and environmental challenges, paving the way for innovative applications in bioengineering, bioremediation, and sustainable soil management.

## Methods

### Prosser soil viral AVG identification

We previously recovered soil AVGs with a broad range of predicted metabolic functions from 2.5 Tb metagenomes sequenced from a Warden silt loam marginal grassland soil in Prosser, Washington (46°15′04″N, 119°43′43″W)^7,28,29^. Sequence similarity of the 37 putative AVG-encoded proteins was assessed against the NCBI Reference Sequence Database (RefSeq; https://www.ncbi.nlm.nih.gov/refseq/) and RCSB Protein Data Bank (PDB; https://www.rcsb.org/) databases using blastp (protein-protein BLAST; blast.ncbi.nlm.nih.gov/Blast.cgi?PAGE=Proteins). For each AVG-encoded protein, the maximal sequence identity and fractional coverage against the best hit in each database are reported in **Extended Data 1**. Structure predictions for each AVG were generated using AlphaFold2 (v2.2.0). The predicted AlphaFold2 structures for V-C12DO were compared to experimental structures in the PDB or determined in this work using the Dali webserver^45^ (http://ekhidna2.biocenter.helsinki.fi/dali/).

### Detection of putative transcription of the soil viral AVGs in metatranscriptomes of the Prosser site

AVGs with consistent functional annotations as their close hits in NCBI RefSeq and RCSB PDB databases were screened for their putative expression within the soil microbiomes. The quality-filtered forward reads of the 62 published metatranscriptomes sequenced from soils of the same field site^30,32^ were indexed and mapped to the AVGs using bwa-mem2 (v2.2.1). Samtools (v1.18) was used to sort and screen for the mapped reads (‘-F 0×4’). The mapped reads were further extracted and searched against the 5.4 M dereplicated genes previously predicted from the non-viral contigs assembled from the same metagenomes where the AVGs were detected^7^. The non-viral genes were dereplicated using Mmseqs (v13.4511) with default parameters (–e 0.001 –-min-seq-id 0.95 –c 0.90). The metatranscriptomic read mappings that were not specific to viral AVGs contained potential false positives and thus were excluded from the AVG transcript counts. The mapped reads for each AVG were counted, normalized to Reads Per Kilobase per Million mapped reads (i.e., RPKM), and used to conservatively estimate the putative transcription of these AVGs. We acknowledge that the screening criteria can underestimate the putative transcription of the soil viral AVGs sharing high sequence similarity with host genes. Therefore, this step was not to quantify the AVG transcripts but to conservatively screen the AVGs whose transcription was detectable in the soil microbiome.

### AVG product screening for function testing

To examine the enzymatic activity of the transcriptionally active AVG, we constructed *E. coli* expression plasmids for the AVGs (see below for details) whose predicted protein products were most suitable for enzymatic assays based on their inferred catalytic functions. These constructs were evaluated for large-scale soluble protein expression using autoinduction methods (500 mL cultures). One construct yielded a positive result, defined by the presence of the target protein as the predominant band on a size exclusion column (Superdex75 or Superdex200) following metal-affinity chromatography (Nickel-NTA). The successful target was predicted to be a catechol 1,2-dioxygenase (C12DO), which was subsequently subjected to functional characterization to elucidate its potential role in the soil microbiome.

### Sequence embedding and distance-based analysis of C12DO diversity

To quantify global sequence-level divergence among catechol 1,2-dioxygenases, we automatically retrieved protein sequences from the NCBI, PDB, and UniProt databases (accessed December 2025), together with associated taxonomic lineage information and database-labeled oligomeric state, using a customized Python workflow. Retrieved sequences were dereplicated using CD-HIT (v4.8.1), and C12DO annotations were verified using the KEGG ortholog K03381 Hidden Markov Model (HMM) with kofamscan (v1.3.0; maximum e-value 8.00 × 10^−^¹□, minimum bit score 54.6).

A total of 10,950 dereplicated protein sequences spanning domains of life were embedded using the ESM-2 protein language model^66^ (esm2_t6_8M_UR50D), which generates numerical representations that capture contextual sequence features learned from large-scale protein sequence data. Embeddings were computed on the CPU in batches, producing one fixed-length embedding vector per full-length protein sequence. Pairwise sequence divergence was quantified using cosine distance in the embedding space. To visualize the global structure of the C12DO sequence diversity, embedding vectors were projected into two dimensions using Uniform Manifold Approximation and Projection (UMAP) with default parameters. The viral catechol 1,2-dioxygenase (V-C12DO) was identified by its sequence identifier and used as a reference point for visualizing nearest neighbors and for downstream distance-based analyses.

For statistical analysis, the cosine distance between each sequence and V-C12DO was treated as a continuous response variable. Sequence metadata, including sequence length, domain, and taxonomic lineage, were merged with the distance table. The relative contributions of sequence length, domain, and lineage to variation in embedding distance were evaluated using analysis of variance (ANOVA). Effect sizes were quantified using partial eta-squared, and the proportion of the total sum of squares explained by each factor was calculated to assess their relative importance.

### HMM-guided alignment and length-independent residue conservation analyses

To assess residue-level conservation in a coordinate system that is robust to length variation and truncation, all C12DO sequences were aligned to the K03381 HMM using hmmalign (HMMER v3.3.2). We analyzed residues in HMM match space, which captures model-defined core positions and provides a length-independent indexing framework for comparing homologs across domains of life.

Key catalytic residues, specifically the two tyrosines (Y) and two histidines (H) that coordinate the active-site iron, were first specified relative to ungapped reference sequences. These residues were mapped to alignment columns by tracking ungapped residue positions through the HMM-guided alignment, and then converted to HMM match column indices. This mapping links biochemically defined sites to a shared, length-independent coordinate system, enabling direct comparisons even when sequences contain insertions, deletions, or N– or C-terminal truncations.

For each HMM match column, residue composition was computed across all aligned sequences. Gaps and ambiguous residues were excluded from residue counts. For each column, we quantified the total number of sequences, the number of valid (non-gap, non-ambiguous) residues, and per-residue frequencies. These statistics were computed both globally across all sequences and within lineage-resolved groupings. All calculations and downstream visualizations were indexed by HMM match columns to maintain consistent positional comparability across sequences with differing lengths and degrees of truncation.

### Primary amino acid sequence and tertiary structure alignments

To further investigate the functional divergence within the cluster containing bacterial representatives, we conducted detailed sequence and structural comparisons. Multiple sequence alignment of the primary amino acid sequence for V-C12DO and bacterial C12DOs, comprising both dimeric and monomeric forms with available crystal structures in the metal-bound state, was obtained using the Clustal Omega MSA webserver (https://www.ebi.ac.uk/jdispatcher/msa/clustalo). Structural alignments were visually inspected and compared using PyMOL (v 2.5.7).

### Expression and purification of a phage catechol 1,2 dioxygenase

The gene encoding V-C12DO was synthesized with a 21-residue N-terminus extension (MGSS<u>HHHHHH</u>SSGENLYFQGH-) containing a poly-histidine metal affinity tag (underlined) and a Tobacco Etch Virus (TEV) protease cleavage site (bold) and inserted into a pET21 plasmid. The plasmid was used to transform chemically competent *E. coli* BL21(DE3) (Novagen, Darmstadt, Germany) from which multiple (5-7), ∼1 mL, ∼15% glycerol, frozen (−80 °C) stocks (LB media, OD_600nm_ = ∼0.8) were prepared from a single colony picked off a streaked LB agar plate. A single ∼1 mL stock was used to seed 20 mL of LB medium that was grown to an OD_600nm_ of ∼ 0.8 and then transferred to 500 mL of autoinduction ZY medium^67^ (2 L flasks, 180 rpm shaker, 0.100 ug/uL ampicillin, 37 °C). Upon reaching an OD_600nm_ of approximately 1, the temperature was lowered to 20 °C and the media was spiked with 500 μL of 10 mM FeCl_3_. The cells were harvested by gentle centrifugation the next day (∼16 h later) and then frozen (−80 °C). The frozen pellet from a 500 mL culture was later thawed by resuspension in ∼ 32 mL of NTA wash buffer (50 mM sodium phosphate, 300 mM NaCl, 10 mM imidazole, pH 7.8), followed by sonication (2 min) before and after three passes through a French Press (SLM Aminco, Rochester, NY). Following centrifugation, the decanted soluble fraction was sonicated for 1 min and the protein purified using a conventional two-step purification protocol: metal chelate affinity chromatography on a 20 mL Ni-Agarose 6 FastFlow column (GE Healthcare, Piscataway, NJ) followed by gel-filtration chromatography on a Superdex75 HiLoad 26/60 column (GE Healthcare, Piscataway, NJ). Fractions containing the target protein after the last column step were pooled and concentrated to ∼ 5 mg/mL (Protein Buffer: 100 mM NaCl, 20 mM Tris, 1 mM DTT, pH 7) and stored at 4 °C until performing the experiments. Yields of V-C12DO were between 5-10 mg per liter ZY medium.

To assist the characterization of the oligomeric state of V-C12DO, a glycerol stock of an expression host containing the recombinant plasmid for a C12DO gene from *Burkholderia multivorans* (*Bm*-C12DO) was obtained from the Seattle Structural Genomics Center for Infectious Diseases (SSGCID-ID: BumuA.00117.a; UniProt: A0A0H3KXJ8_BURM1)^68^. Iron-bound *Bm*-C12DO crystallized as a dimer (9DR8) and the PDBe PISA (https://www.ebi.ac.uk/pdbe/pisa/) analysis of the crystal structure reports that this interface results in a stable dimeric quaternary structure in solution. Iron-bound, ^15^N-labelled *Bm*-C12DO and unlabeled *Bm*-C12DO were prepared in the same Protein Buffer following the strategy described for V-C12DO using auto-induction with minimal media and ZY media, respectively. Yields of purified *Bm*-C12DO were in the 20-40 mg/L range. In pursuit of a V-C12DO crystal, a construct in pET-22b(+) was prepared by removing the N-terminal purification tag and adding a shorter, 8-residue, purification tag (−LEHHHHHH) at the C-terminal (V-C12DO*; SSGCID-ID: MesoA.00117.a, Genescript, Piscataway, NJ). Yields of purified V-C12DO* were more than 10-fold greater than for V-C12DO. Nitrogen-15 labelled V-C12DO* samples were prepared with 500 mL of autoinduction minimal media spiked with 500 μL of 10 mM FeCl_3_ prior to lowering the temperature to 20 °C. The iron-bound C12DO samples expressed in the presence of excess iron were colored red/purple.

### Crystallization, X-ray data collection, and structure solution

Numerous attempts to obtain crystals for V-C12DO, including the use of many commercial screening kits and the services of the Hauptman-Woodward High-Throughput Crystallization Screening Center (1536 conditions)^69^, proved unsuccessful. Proteolytic removal of the short, N-terminal 21-residue purification tag on V-C12DO also did not yield crystals that diffracted well. Suitable crystals were only obtained by replacing the V-C12DO 21-residue N-terminal purification tag with a C-terminal, uncleavable, 8-residue tag, –LEHHHHHH (V-C12DO*). This construct resulted in crystals that were grown using the hanging drop method by mixing 1.6 uL of protein (∼ 7 mg/mL in Protein Buffer) and 1.6 uL of precipitant (Anatrace (Maumee, OH) Top96 condition #44 (0.1 M Bis-TrisHCl, pH 6.5, 20% (w/v) PEG MME 5000)). Crystals, buried in protein precipitate and sensitive to protein concentration, first appeared 2-3 days following mixing and incubation at room temperature. X-ray diffraction data were collected at a wavelength of 0.9786 Å and temperature of 100 K at the National Synchrotron Light Source-II (NSLS-II) NYX beamline 19-ID using an Eiger2 XE 9M pixel array detector. Intensities were integrated using XDS (v20210323)^70^ via Autoproc (v1.0.5)^71^ and the Laue class analysis and data scaling were performed with Aimless (v0.8.2)^72^. Structure solution was conducted by molecular replacement with Phaser (v2.8.3)^73^ using a model generated with the AlphaFold3 server as the search model. Refinement and manual model building were conducted with Phenix (v1.21.2-5533)^74^ and Coot (v0.9.8.96)^75^, respectively. The final structure had a Molprobity score of 1.23 (98^th^ percentile) with 98.5% and 1.5% of the backbone dihedral angles in preferred and allowable Ramachandran regions, respectively. Relevant crystallographic data are provided in **Table 2**. The PDB was searched for structures similar to V-C12DO with DALI (http://ekhidna2.biocenter.helsinki.fi/dali/).

### Standard enzyme assay and kinetic parameter determination with catechol

The activity of V-C12DO for catechol (Sigma-Aldrich, St. Louis, MO) was measured spectrophotometrically at 260_nm_ using a standard curve prepared for the product, *cis,cis*-muconic acid (Sigma-Aldrich, St. Louis, MO) (5-50 mM), in our standard reaction buffer (50 mM sodium phosphate, 100 mM NaCl, pH 6.5). Michaelis-Menton enzyme catalytic parameters (K_m_ and V_max_) for V-C12DO were calculated using a Lineweaver-Burk analyses by measuring the initial linear rate of product formation over the first 30 s after the addition of different amounts of catechol (1 to 200 μM; volumes ≤ 10 μL from 1, 10, or 100 mM catechol stock solutions prepared in water) to a fixed amount of enzyme (0.2 μM, 2 μL of a 0.1 mM stock solution of V-C12DO). The reactions were performed at room temperature (20 °C) in a quartz cell with a 1 cm path length using 1 mL of the standard reaction buffer. A Nanodrop 2000c (Thermo Fisher Scientific, Waltham, MA) was used for all spectrophotometric measurements.

### Determination of substrate specificity

To explore substrate specificity, the following catechol derivatives were assayed as substrates for V-C12DO using the standard reaction conditions: 3-methylcatechol (2-methylmuconic acid; e_260_ = 18,000 M^−1^cm^−1^), 4-methylcatechol (3-methylmuconic acid; e_255_ = 14,300 M^−1^cm^−1^), 4-ethylcatechol (3-ethylmuconic acid; e_255_ = 14,300 M^−1^cm^−1^), 4-chlorocatechol (3-choloromuconic acid; e_259_ = 12,500 M^−1^cm^−1^), 4,5-dichlorocatechol (3,4-dichloromuconic acid), and pyrogallol (2-hydroxymuconic acid (e_296_ = 14,500 M^−1^cm^−1^)^76^, and the methods in the previous paragraph. These substrates were purchased from Sigma-Aldrich (St. Louis, MO). Note that a molar extinction coefficient for 3-ethylmuconic acid could not be found, and therefore, the value for 3-ethylmuconic acid was used in the calculations. All experiments were performed at 20 °C, in triplicate, with the relative activity referenced against catechol.

### Effects of pH, salinity, and temperature

The optimum pH for V-C12DO activity was determined in solutions containing 100 mM NaCl and the following buffers: 50 mM sodium acetate (pH 2.8, 4.5, and 5.7), 50 mM BisTris (pH 6.0), 50 mM sodium phosphate (pH 6.5, 7.0, and 7.5), 50 mM Tris (pH 8.4 and 9.0), and 50 mM CAPS (pH 10.1). Observing that the maximum activity was with the pH 6.5 buffer (our standard reaction buffer), the experiments to evaluate the impact of salinity and temperature on V-C12DO activity were performed in this buffer by varying the NaCl concentration between 0.1 and 3.0 M, and the temperature between 5 and 60 °C. The reactions were performed as described for the standard enzyme assay, except for starting all the reactions with 100 μM of catechol (10 μL addition of a 10 mM catechol stock solution prepared in water). All experiments were performed at 20 °C, in triplicate, with the relative activity based on the condition with the fastest activity.

### Native mass spectrometry

Samples were buffer exchanged into 0.2 M ammonium acetate using BioRad micro Bio-Spin 6 columns. They were then loaded into 0.78 I.D. glass capillary emitters that were pulled in-house and analyzed using a Waters Synapt G2-Si mass spectrometer in sensitivity mode using the following instrument settings: 1.0-1.5 V spray voltage, mass range 400-8000 m/z, trap gas 4 mL/min, source temperature 60 °C, sampling cone 150, source offset 60, in positive ion mode. The data were deconvolved using UniDec software (v8.0.0)^77^.

### Solution Nuclear Magnetic Resonance spectroscopy

The NMR data were collected at 20 °C on ^15^N-labelled samples (∼1 mM) in a buffered solution (100 mM NaCl, 20 mM Tris, 1 mM DTT, pH 7) using a Varian NMR spectrometer operating at a ^1^H resonance frequency of 600 MHz equipped with an HCN-cryoprobe and pulse field gradients. The ratio of collective backbone amide ^15^N T_1_ and T_1rho_ measurements obtained using an adapted ^1^H-^15^N HSQC experiment to collect ^15^N-edited one-dimensional proton spectra was used to estimate an overall rotational correlation time, t_c_, for V-C12DO* and *Bm*-C12DO at 20 °C.

## Data availability

The sequencing data (metagenome and metatranscriptome) used in the study have been deposited along with our previous publications (metagenome^28,29^; metatranscriptome^30,32^). Coordinates and structure factors for the V-C12DO structure have been deposited to the Worldwide Protein Databank (wwPDB) with the accession code 9YH9. Input files used to generate both main and supplementary figures are deposited as Source Data at Zenodo^78^.

## Code availability

Scripts for the sequence analyses are available at a public GitHub repository, https://github.com/Ruonan0101/C12DO_MS/tree/main.

## Acknowledgements

This program is supported by the U. S. Department of Energy, Office of Biological and Environmental Research, Genomic Sciences Program under FWP 70880. PNNL is a multi-program National Laboratory operated by Battelle for the DOE under Contract DE-AC05-76RLO 1830. A portion of this work was supported by the Seattle Structural Genomics Center for Infectious Diseases (SSGCID), funded by the National Institute of Allergy and Infectious Diseases, National Institutes of Health, Department of Health and Human Services, under Contract No.: 75N93022C00036. This research used resources of the NYX beamline 19-ID, supported by the New York Structural Biology Center, at the National Synchrotron Light Source II, a U.S. Department of Energy (DOE) Office of Science User Facility operated for the DOE Office of Science by Brookhaven National Laboratory under Contract No. DE-SC0012704. The NYX detector instrumentation was supported by grant S10OD030394 through the Office of the Director of the National Institutes of Health.

## Ethics declarations

Competing interests

The authors declare no competing interests.

## Extended data

**Extended Data 1. AVGs identified in the studied site.**

**Extended Data 2. Residue conservation profile across the HMM-anchored alignment.** Residue fractions are shown for each aligned HMM column across the global C12DO alignment, plotted in three windows spanning columns 102 to 413. Points are colored by residue identity at each column and are transparent when the residue fraction at that position is < 50%. Residues are labeled in columns where the maximum residue fraction is ≥ 50%. The four known key-residue columns are marked by red dashed lines, and fully conserved columns (maximum fraction = 100%) are marked by grey dashed lines.

**Extended Data 3: Crystallographic data for V-C12DO\***.

## Source data

Source Data for Figure 1

Source Data for Figure 2

Source Data for Figure 3

Source Data for Figure 4

Source Data for Figure 5

Source Data for Extended Data 2

