## Supplementary figures and images for "A truncated soil phage catechol 1,2-dioxygenase illustrates how viruses preserve and disseminate auxiliary catalytic functions in the soil microbiome"

### Extended Data 2

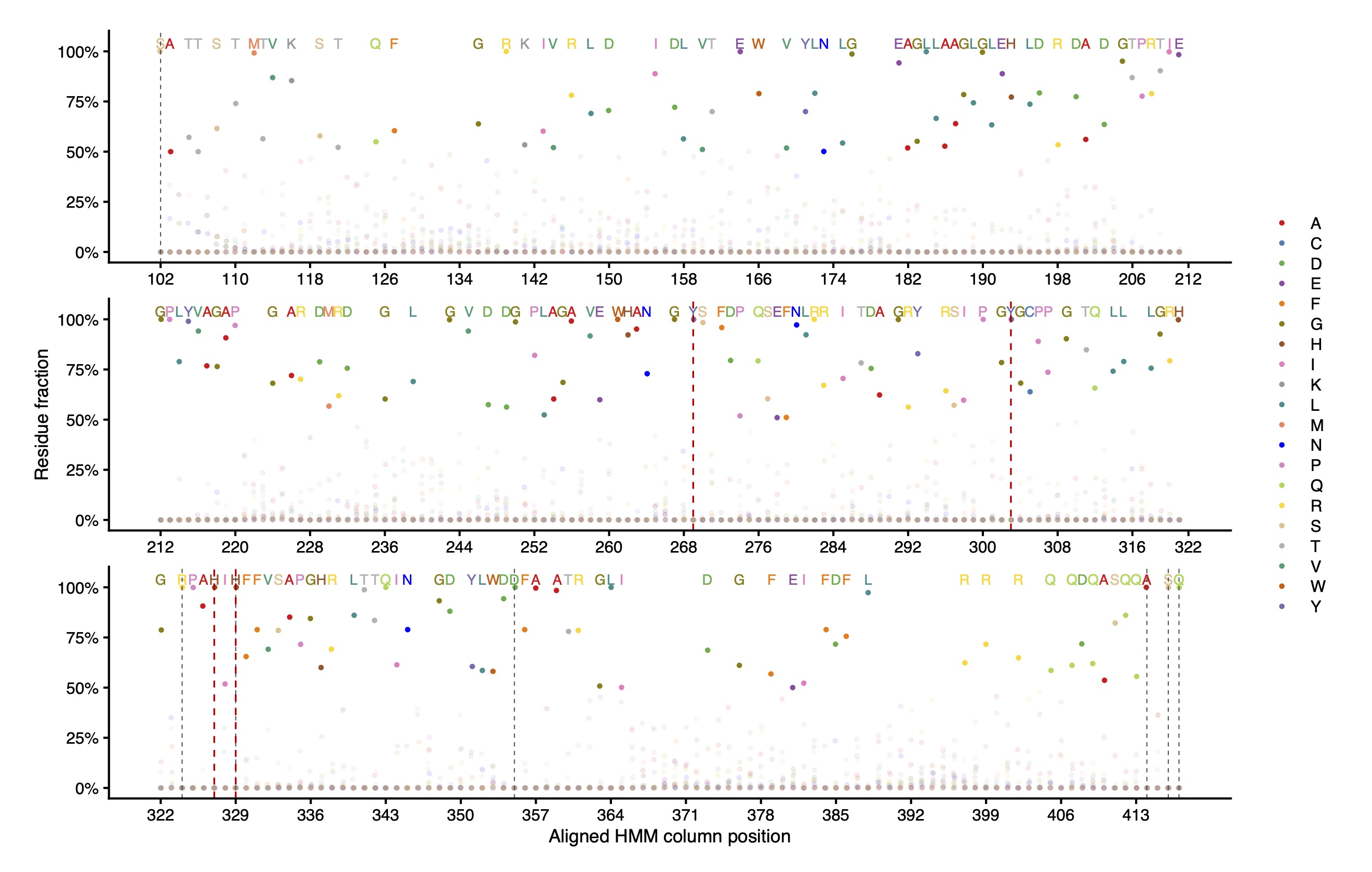

### Supplementary Figure 1

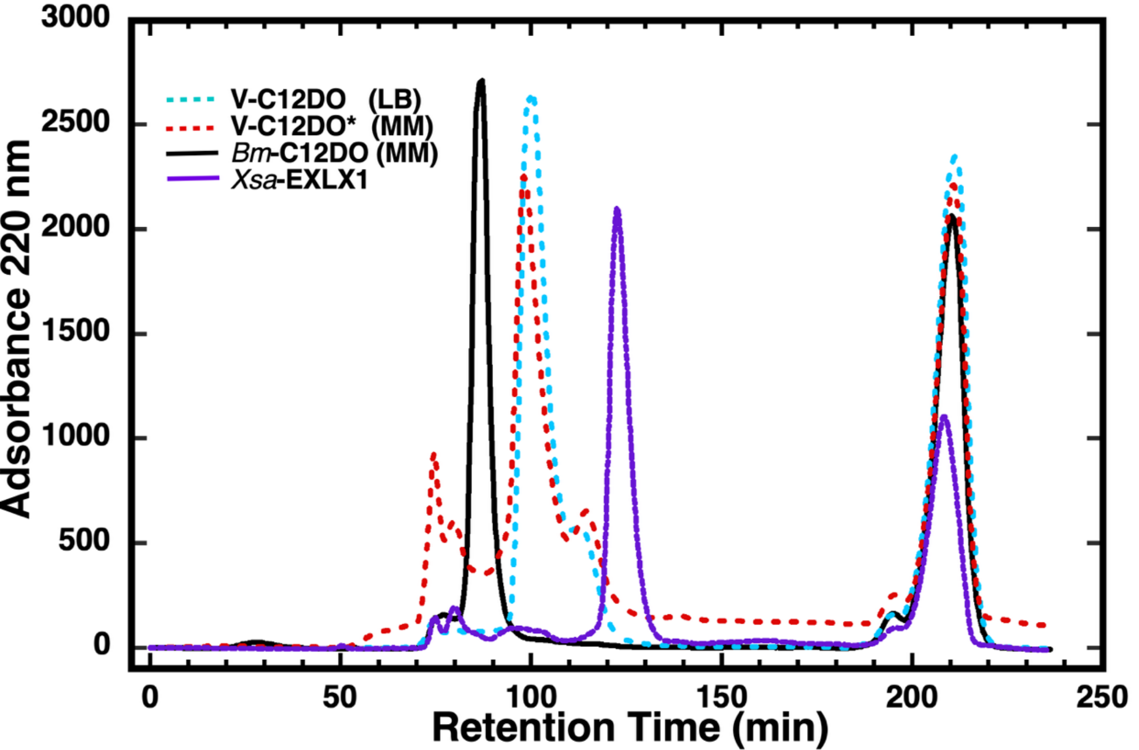

### Supplementary Figure 2

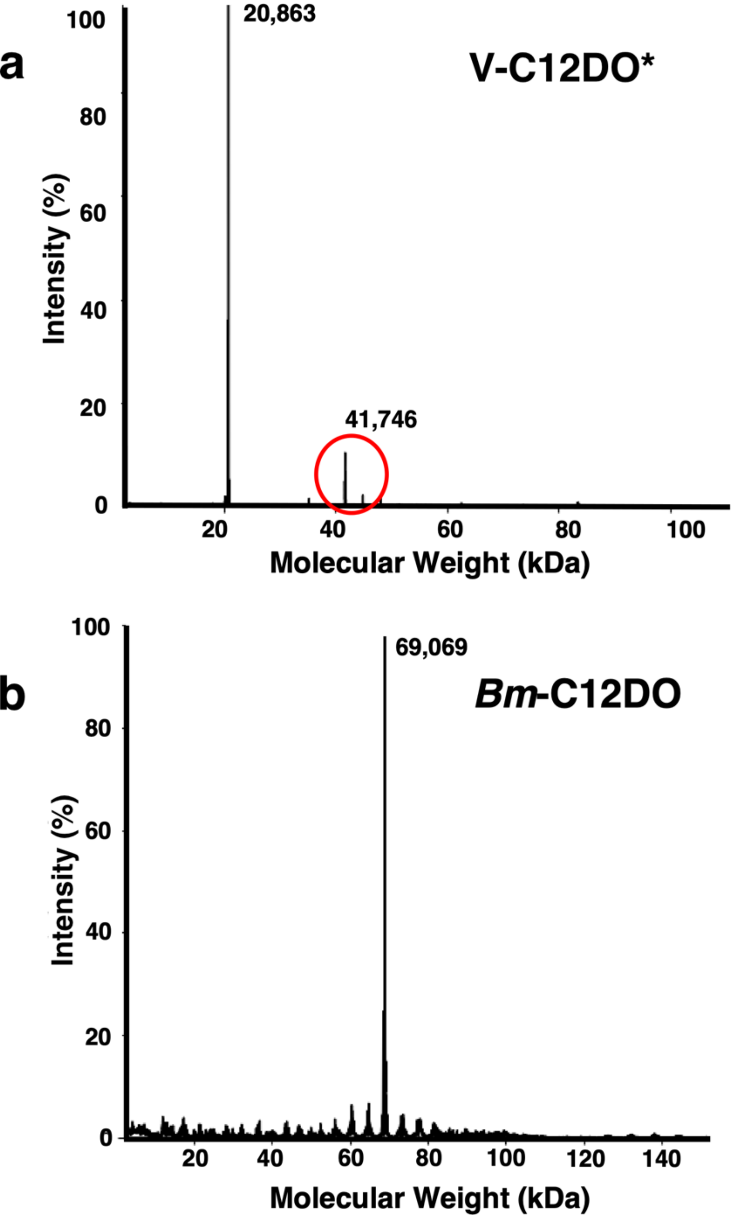

### Supplementary Figure 3

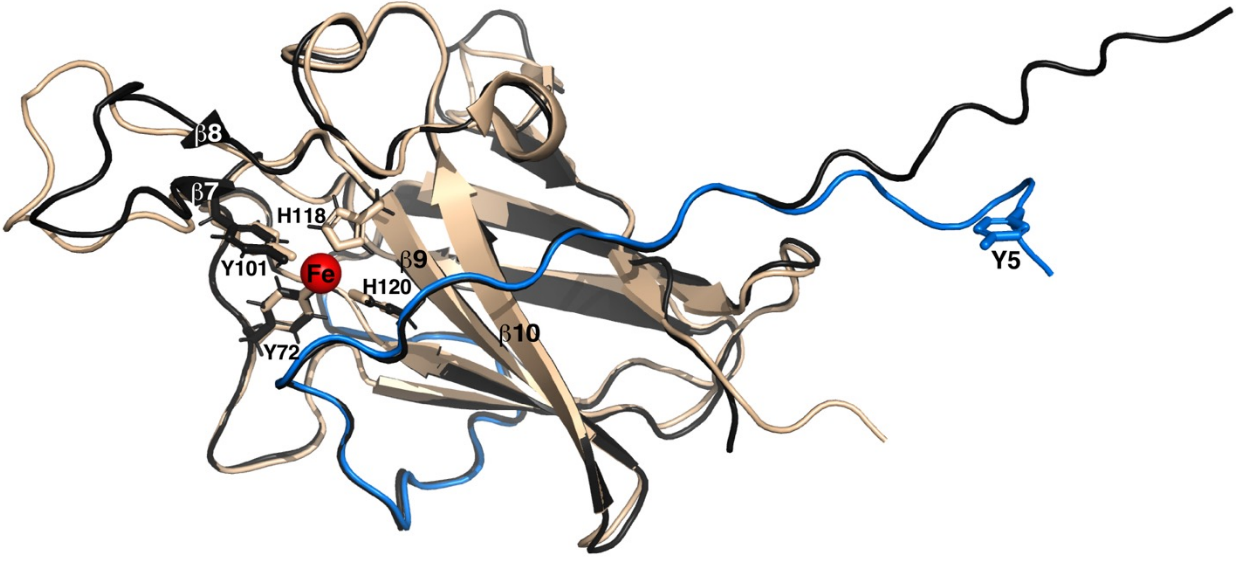
