## Extended Data 3 for "A truncated soil phage catechol 1,2-dioxygenase illustrates how viruses preserve and disseminate auxiliary catalytic functions in the soil microbiome"

**Table 1** **Data collection and refinement statistics for V-C12DO***

|  | V-C12DO* |
| --- | --- |
| **Data collection** |  |
| Space group | *P*2_1_2_1_2_1_ |
| Cell dimensions |  |
| *a*, *b*, *c* (Å) | 47.743, 56.46, 138.46 |
| α, β, γ (°) | 90, 90, 90 |
| Resolution (Å)^1^ | 47.74-1.65 |
| *R*_merge_ | 11.6 (194.0) |
| *I* / σ*I* | 15.5 (1.7) |
| Completeness (%) | 100 (100) |
| Redundancy | 13.1 (13.9) |
| **Refinement** |  |
| Resolution (Å) | 45.14-1.65 |
| No. reflections (working/test) | 43,593 / 2,325 |
| *R*_work_ / *R*_free_ |  |
| No. atoms |  |
| Protein | 2739 |
| Iron | 2 |
| Water | 254 |
| *B*-factors |  |
| Protein | 33.3 |
| Iron | 25.8 |
| Water | 34.9 |
| R.m.s. deviations |  |
| Bond lengths (Å) | 0.009 |
| Bond angles (°) | 1.004 |

^1^One xtal used for data collection with the values in parentheses for the highest-resolution shell.
